# Variation and selection on codon usage bias across an entire subphylum

**DOI:** 10.1101/608042

**Authors:** Abigail L. Labella, Dana A. Opulente, Jacob L. Steenwyk, Chris Todd Hittinger, Antonis Rokas

## Abstract

Variation in synonymous codon usage is abundant across multiple levels of organization: between codons of an amino acid, between genes in a genome, and between genomes of different species. It is now well understood that variation in synonymous codon usage is influenced by mutational bias coupled with both natural selection for translational efficiency and genetic drift, but how these processes shape patterns of codon usage bias across entire lineages remains unexplored. To address this question, we used a rich genomic data set of 327 species that covers nearly one third of the known biodiversity of the budding yeast subphylum Saccharomycotina. We found that, while genome-wide relative synonymous codon usage (RSCU) for all codons was highly correlated with the GC content of the third codon position (GC3), the usage of codons for the amino acids proline, arginine, and glycine was inconsistent with the neutral expectation where mutational bias coupled with genetic drift drive codon usage. Examination between genes’ effective numbers of codons and their GC3 contents in individual genomes revealed that nearly a quarter of genes (381,174/1,683,203; 23%), as well as most genomes (308/327; 94%), significantly deviate from the neutral expectation. Finally, by evaluating the imprint of translational selection on codon usage, measured as the degree to which genes’ adaptiveness to the tRNA pool were correlated with selective pressure, we show that translational selection is widespread in budding yeast genomes (264/327; 81%). These results suggest that the contribution of translational selection and drift to patterns of synonymous codon usage across budding yeasts varies across codons, genes, and genomes; whereas drift is the primary driver of global codon usage across the subphylum, the codon bias of large numbers of genes in the majority of genomes is influenced by translational selection.

**Lay Summary / Significance statement:** Synonymous mutations in genes have no effect on the encoded proteins and were once thought to be evolutionarily neutral. By examining codon usage bias across codons, genes, and genomes of 327 species in the budding yeast subphylum, we show that synonymous codon usage is shaped by both neutral processes and selection for translational efficiency. Specifically, whereas codon usage bias for most codons appears to be strongly associated with mutational bias and largely driven by genetic drift across the entire subphylum, patterns of codon usage bias in a few codons, as well as in many genes in nearly all genomes of budding yeasts, deviate from neutral expectations. Rather, the synonymous codons used within genes in most budding yeast genomes are adapted to the tRNAs present within each genome, a result most likely due to translational selection that optimizes codons to match the tRNAs. Our results suggest that patterns of codon usage bias in budding yeasts, and perhaps more broadly in fungi and other microbial eukaryotes, are shaped by both neutral and selective processes.

## Introduction

One of the first insights drawn from DNA sequence analyses was that synonymous codons are used both non-randomly and in taxon-specific patterns (Air et al. 1976; Fiers et al. 1976; Grantham et al. 1981). These results were surprising given that synonymous codon changes do not alter primary protein structure (i.e., they are silent) and were therefore previously assumed to be selectively neutral. Two major explanations have been put forth to account for the non-random variation in codon usage seen within and across species, namely natural selection and neutral processes, such as mutational bias coupled with genetic drift.

The discovery that codon usage is correlated with both the abundance of transfer RNA molecules in the genome and with gene expression levels raised the hypothesis that optimization of codons to match the available tRNA pool (or tRNAome) promotes or regulates translation and suggested a key role for codon usage in translational dynamics (Post et al. 1979; Nakamura et al. 1980; Ikemura 1981a; Ikemura 1981b; Gouy and Gautier 1982; Sharp and Li 1986; Thomas et al. 1988). It is now well established that codon usage influences multiple cellular processes, especially translation. For example, usage of codons corresponding to the tRNA pool, known as codon optimization, has been linked to increased translation speed (Bulmer 1991; Xia 1998; Chevance et al. 2014; Presnyak et al. 2015), accurate tRNA pairing (Stoletzki and Eyre-Walker 2007; Zhou et al. 2009), suppressed premature cleavage and polyadenylation of transcripts (Zhou et al. 2018), and mRNA stability (Presnyak et al. 2015; Radhakrishnan et al. 2016). Conversely, non-optimal codon usage has been associated with translation initiation (Tuller et al. 2010), accurate protein folding (Zhou et al. 2013; Yu et al. 2015; Buhr et al. 2016), and signal recognition particle detection (Pechmann et al. 2014). These molecular discoveries are complemented by a plethora of examples where specific synonymous substitutions have substantial fitness (Agashe et al. 2013; Fragata et al. 2018; Mittal et al. 2018; Ballard et al. 2019) and phenotypic effects in organisms across the tree of life, including *Escherichia coli* (Krisko et al. 2014), *Saccharomyces cerevisiae* (Kliman et al. 2003; She and Jarosz 2018), *Drosophila melanogaster* (Carlini and Stephan 2003), and humans (Chamary et al. 2006; Sauna and Kimchi-Sarfaty 2011; Supek et al. 2014). In summary, there is now substantial evidence to suggest that codon usage bias of certain codons in certain species is under strong selection—often through translational mechanisms.

In the absence of selection or in populations where genetic drift is more powerful than selection, patterns of codon usage bias will reflect the effects of genome-wide mutational pressures, such as mutational bias or GC-biased gene conversion (Sharp and Li 1987; Knight et al. 2001; Chen et al. 2004; Palidwor et al. 2010; Galtier et al. 2018). This was first suspected for species with extreme GC composition biases, such as the Gram positive bacterium *Mycoplasma capricolum*, which has a genomic GC composition of 25%, and only 2% of its codons end with G or C (Sharp et al. 1993). For species like *M. capricolum*, it was hypothesized that biased genome-wide mutational processes, such as mutational bias towards A/T bases and GC-biased gene conversion, would drive patterns of codon usage bias. GC-biased gene conversion has been shown to influence the GC content of third codon positions in an evolutionarily neutral manner in mammals, as well as at recombination hotspots in yeasts (Galtier et al. 2001; Harrison and Charlesworth 2011). Mutational bias has been proposed as the major driver of codon usage bias in diverse studies in a variety of lineages, including bacteria, archaea, plants, and animals (Chen et al. 2004; Wan et al. 2004; Palidwor et al. 2010; Clement et al. 2017). Even in the presence of selection on synonymous codon sites, it has been proposed that background substitution drives codon preference in organisms with widely different GC compositions (Sun et al. 2017). Thus, major differences in codon usage patterns between species are often considered to be primarily driven by neutral mutational changes in GC content (Knight et al. 2001; Chen et al. 2004).

Selective and neutral explanations of codon usage bias are not mutually exclusive, and pioneers in this field were quick to suggest that codon bias is due to a balance between neutral and selective processes (Ikemura 1985; Shields and Sharp 1987; Sharp et al. 1993). It is unclear, however, what that balance is, how it varies across levels of biological organization (e.g., codons, genes, genomes) and across lineages, and what factors influence the balance (Bulmer 1991; Sharp et al. 1993; Sharp et al. 1995; Knight et al. 2001; Hershberg and Petrov 2008; Palidwor et al. 2010).

Budding yeasts (subphylum Saccharomycotina, phylum Ascomycota) present a unique opportunity to examine the impact of neutral and selective processes on codon usage bias for several reasons. First, genomes and genome annotations of 332 species across the subphylum recently became available (Shen et al. 2018), providing a state-of-the-art data set for the study of codon usage bias. Second, the genomic diversity across budding yeasts is comparable to the divergence between different animal phyla or between *Arabidopsis* and green algae, offering us the opportunity to examine variation in patterns of codon usage bias across a highly diverse lineage. Third, budding yeasts exhibit genetic code diversity and are the only known lineage with nuclear codon reassignments. Specifically, three different clades of buddying yeasts have undergone a reassignment of the CUG codon from leucine to serine (two clades) or alanine (one clade) (Kawaguchi et al. 1989; Miranda et al. 2006; Muhlhausen et al. 2016; Riley et al. 2016; Krassowski et al. 2018). Codon reassignments in the Saccharomycotina provide both a challenge and an opportunity in comparing codon usage bias across the subphylum. Finally, for the majority of budding yeast species in our data set we also have metabolic trait (285 species) and isolation environment (174 species) information, which not only illustrates the ecological diversity of this group but allows us to test for other contributors to codon usage bias (Kurtzman et al. 2011; Opulente et al. 2018).

To examine codon usage bias at the codon, gene, and genome levels, we examined the genomes of 327 budding yeast species in the subphylum Saccharomycotina. Analysis of codon usage bias, measured by relative synonymous codon usage (RSCU) revealed diversity in usage at all three levels (codon, gene, genome) examined. This variation in RSCU was highly correlated with GC composition when assessed broadly across the subphylum. Furthermore, the relationship between the relative frequency of each codon and the GC composition of the 3^rd^ codon position showed very small deviations from the neutral expectation, except for codons for three amino acids (proline, arginine, and glycine). However, at the gene level, nearly a quarter of all genes surveyed (381,174/1,683,203; 23%) did not fit the neutral expectation of the relationship between the effective number of codons and synonymous GC composition. In 94% (308/327) of the budding yeast genomes, the overall fit of genes to the neutral expectation was very low. Investigation of possible causes of this deviation revealed that 81% (264/ 327) of budding yeast genomes exhibited moderate-to-high levels translational selection on codon usage bias. While there was no significant correlation between the total number of metabolic traits or isolation environments and selection, the strength of selection was significantly correlated with genomic tRNA gene content (tRNAome). These results suggest that translational selection on codon bias is widespread, but not ubiquitous, in the budding yeast subphylum. Our inference of strong translational selection on codon usage bias suggests that translational regulation has played a major role in the evolution of this group.

## Methods

### Sequence Data

Genomic sequence and annotation data were obtained from a recent comparative genomic study of 332 budding yeast genomes (Shen et al. 2018) (Supplementary Table 1). Genomes of five species from the CUG-Alanine clade were removed from this analysis as their codon reassignment was discovered recently (Muhlhausen et al. 2016; Riley et al. 2016) and could not be accounted for by any existing software. To remove mitochondrial genome sequences from the remaining 327 budding yeast genomes, we employed blastn, version 2.6.0+ (Altschul et al. 1990; Camacho et al. 2009) with 56 partial or complete Saccharomycotina mitochondrial genomes (Supplementary Table 2) as our input queries. Hits that had 30 percent or more sequence identity to mitochondrial sequences were removed from our analyses. Similarly, protein-coding gene sequence data from the 327 genomes were filtered for mitochondrial genes by blasting (blastx) against mitochondrial protein-coding sequence data from 37 Saccharomycotina species (Supplementary Table 3). The coding sequences were further filtered to conform to the required input for the species-specific tRNA adaptation calculations by stAIcalc, version 1.0 (Sabi and Tuller 2014). This filtering step removed all coding sequences that did not begin with the start codon ATG, did not have a whole number of codons, or were shorter than 100 codons (Supplementary Table 1). Codons containing ambiguous bases were also removed.

### Codon usage bias calculations

To examine the variation in codon usage across the yeast subphylum, we calculated the relative synonymous codon usage (RSCU) for each codon in the 1,683,203 protein-coding genes of the 327 budding yeast genomes that remained after filtering. RSCU is the observed frequency of a synonymous codon divided by the frequency expected if all the synonymous codons were used equally (Sharp and Li 1986). We computed RSCU values using DAMBE7, version 7.0.28 (Xia 2018), because it allowed us to accommodate the known nuclear codon reassignment in the CUG-Ser1 and CUG-Ser2 clades (Kawaguchi et al. 1989; Miranda et al. 2006; Muhlhausen et al. 2016; Riley et al. 2016; Krassowski et al. 2018).

To examine broad patterns of codon usage, hierarchical clustering of all RSCU values for each species was calculated and visualized in the R programming environment. To investigate which codons drive between-species differences in codon usage, we performed correspondence analysis of RSCU values (Grantham et al. 1981). This technique is highly suitable and informative because it reduces the high number of dimensions present in codon usage statistics into a very small number of axes (Grantham et al. 1980; Suzuki et al. 2008).

To examine the influence of phylogeny on the observed variation in codon bias, we computed two measures of phylogenetic signal in R, Pagel’s λ (Pagel 1999) and Blomberg’s K (Blomberg et al. 2003). The phylogeny used for this analysis was obtained through maximum likelihood-based inference from a data matrix comprised of 2,408 genes obtained from Shen et al. (2018).

### Mutational bias and codon usage

To assess the role of mutational bias in determining the observed patterns of codon bias in the yeast subphylum, we tested the observed patterns against neutral expectations, both across species and across codons. Between-species patterns in codon usage bias were measured by calculating the Pearson’s correlation of the RSCU of each codon against the GC composition of the 3^rd^ codon position (GC3) across all genes in each genome, for each of the 327 species. To account for the observed phylogenetic dependence within both variables, we also assessed the relationship between RSCU and GC3 using the phylogenetic generalized least squares (PGLS). The influence of mutational bias within each set of codons encoding an amino acid was assessed by comparing the equilibrium solutions for relative codon frequencies based on GC3 content generated by Palidwor et al. (2010) to the empirical values. Observed relative codon frequencies were calculated as the total number of observations of a codon divided by the total number of observations of the corresponding amino acid. Total codon counts within the genomes were calculated in DAMBE version 7.0.28 (Xia 2018). For each codon, predicted values of relative frequency were generated from the corresponding equilibrium solution. R^2^ values were then calculated based on the predicted and empirical relative frequency values. Data from the 98 genomes present in the CUG-Ser1 and CUG-Ser2 clades were removed from the analyses of the amino acids leucine and serine.

To assess the influence of mutational bias within every genome, we compared the effective number of codons (ENC) (Wright 1990) of each gene to the synonymous GC3 proportion of that gene. The NC for each gene within the 327 genomes was computed in DAMBE version 7.0.28 using the improved index created by Sun et al. (2013), which allows for CUG codon reassignments to serine (Xia 2018). This distribution was compared against the predicted neutral distribution proposed by dos Reis et al. (2004) using the suggested parameters. This neutral distribution is a modified version of Wright’s proposed function (Wright 1990) for calculating ENC (dos Reis et al. 2004). We computed an R^2^ value between the observed and empirical ENC values based on the GC3 of each gene. To ensure that R^2^ values were not driven by phylogenetic signal, we calculated Blomberg’s K for the R^2^ values.

### Calculation of selection on codon usage

To determine if selection on translational processes has optimized the codon usage within each species, we tested if there is a significant correlation between the selective pressure on a gene and its level of optimization to the tRNAome for every genome. First, the species-specific value for each codon’s relative adaptiveness (wi) was calculated in stAIcalc, version 1.0 (Sabi and Tuller 2014). Calculation of wi values requires genomic tRNA counts, which we calculated in tRNAscan-SE 2.0 for all species (Lowe and Chan 2016). The results from tRNAscan-SE 2.0 correctly identified the CUG-Ser1 and CUG-Ser2 tRNAs that have a CAG anticodon but the recognition elements for serine (Supplementary Table 4). The species-specific tRNA adaptation index of each gene was then calculated by taking the geometric mean of all wi values for the codons (except the start codon). One drawback of stAIcalc is that it does not account for the nuclear codon reassignment in the CUG-Ser1 and CUG-Ser2 clades. Therefore, we also tested all genomes after removing all CUG codons from all sequences.

To test whether selection has influenced codon usage bias, we calculated the S-value proposed by dos Reis et al. (2004). This metric is the correlation between the tRNA adaptation index (stAI) and the confounded effects of the selection effect of the codon usage of a gene and uncontrollable random factors. Ultimately, the S-value measures the proportion of codon bias variance that cannot be explained by mutational bias or random factors alone. S-values were calculated with the R package tAI.R, version 0.2 (https://github.com/mariodosreis/tai) for each genome using the previously calculated stAI values. We calculated the S-value twice for each genome: once with CUG codons included and once without CUG codons.

To test whether the S-value for a given genome significantly deviated from what would be expected under neutrality, we ran a permutation test. Specifically, we ran 10,000 permutations where each genome’s wi values were randomly assigned to codons, the tAI values were then recalculated for each gene, and the S-test was run on that permutation. A genome’s observed S-value was considered statistically significant if it fell in the top 5% of the distribution formed by the 10,000 values obtained by the permutation analysis.

To investigate which features may influence the level of translational selection occurring within a genome, we tested the contributions of tRNAome size (calculated from tRNA-scan-SE), genome size, number of predicted coding sequences, total number of reported metabolic traits, and total number of reported isolation environments (Shen et al. 2018) on S-value variation. We preformed linear regression analysis on individual and combinations of variables in R. In addition to the linear models, we tested a Gaussian distribution on a subset of features based on visual inspection. We also tested a PGLS analysis on S-value distribution to examine correlations that may be corrected by phylogenetic consideration.

## Results

### Budding yeast genomes exhibit substantial variation in codon usage

To measure variation in codon usage bias across budding yeast genomes, we measured the RSCU of each codon in each Saccharomycotina species. Hierarchical clustering of the codons revealed three major groups of codons (Fig. 1). One group contained codons that were generally overrepresented (RSCU > 1) in budding yeast genomes, which included A/U-ending codons and one G/C-ending codon (UUG). The next group contained mostly G/C-ending codons and two A/U-ending codons (AUA and GUA) that were generally underrepresented (RSCU < 1) across budding yeast genomes. Finally, the smallest group contained A/U-ending codons (CUA, UUA, CGA, GGA, AUA, CCU, and GUA) that were relatively underrepresented across some budding yeast genomes as compared to the first set of A/U-ending codons. Interestingly, the underrepresentation of the CUA codon, which encodes leucine, was driven most strongly by the CUG-Ser1 and CUG-Ser2 clades where the CAG leucine codon has been recoded as serine (Fig. 1).

**Figure 1.**
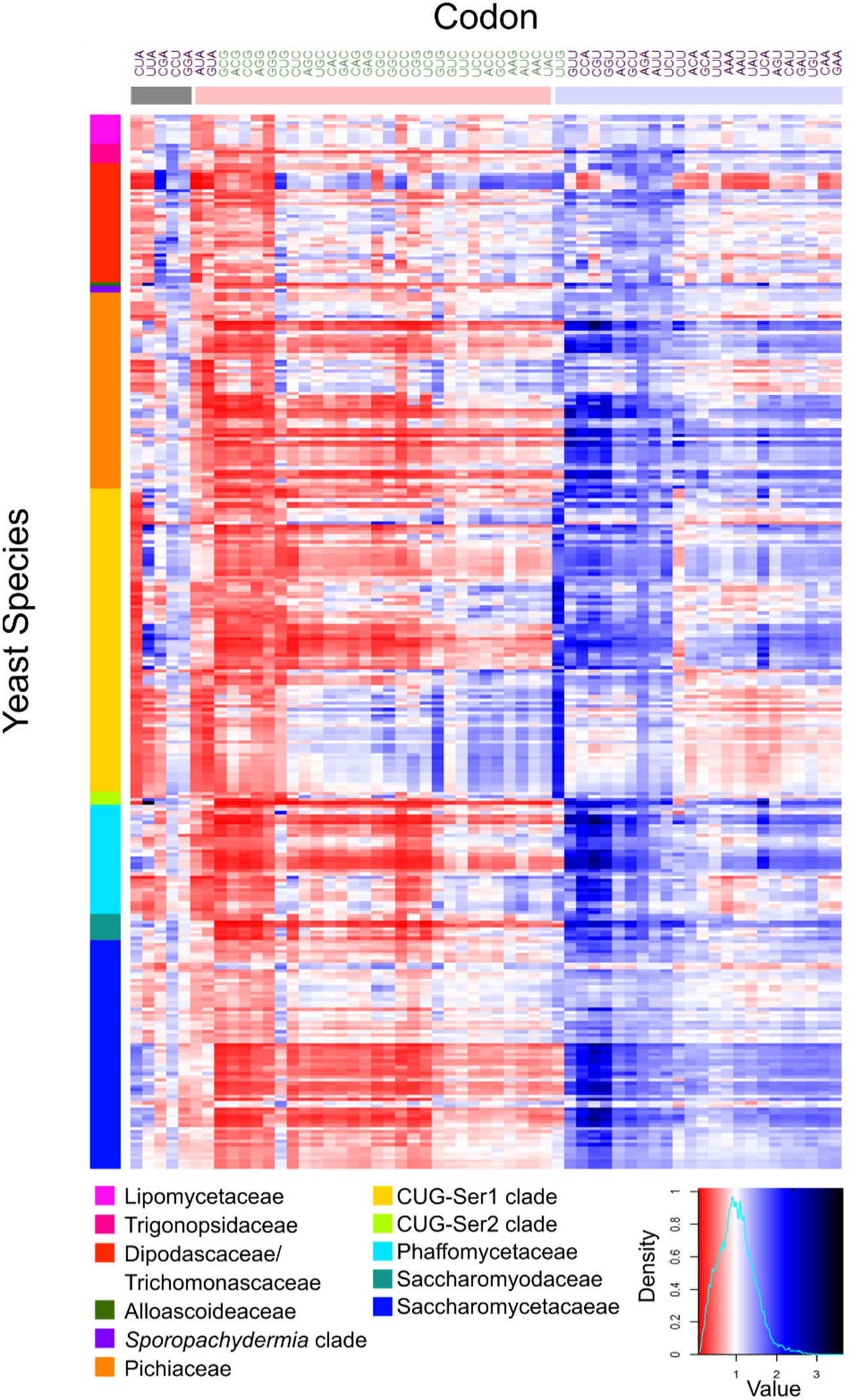
Relative synonymous codon usage (RSCU) analysis revealed an overrepresentation of A/U-ending codons across most of the Saccharomycotina subphylum. Columns correspond to the 59 non-degenerate, non-stop codons; A/U-ending codons are shown in in purple font, and GC-ending codons are shown in green font. Rows correspond to the 327 Saccharomycotina species colored by major clade, following the recent genome-scale phylogeny of the subphylum (Shen et al. 2018). Blue cells indicate overrepresented codons (RSCU > 1) and red cells indicate underrepresented codons (RSCU < 1). Codons were clustered (using hierarchical clustering) by RSCU value into three general groups (shown by horizontal bars of different colors): underrepresented A/U-ending codons (grey bar), underrepresented codons mostly ending in G/C (red bar), and overrepresented codons mostly ending in A/U (blue bar).

### Genome-level variation in codon usage corresponds with mutational bias

To summarize the overall variation in codon usage between species, we conducted a correspondence analysis on RSCU across all 327 species. The majority of the variation in codon usage between species was described by the first dimension of the correspondence analysis (66.891%; Fig. 2), which was driven by differential usage of codons that vary at the third codon position, with the codons UUA, CGU, GGC and GUG making the largest contributions (Supplementary Figure 1a). The second axis, which explained 7.093% of the variation in codon usage, showed some clustering by clade, with the CUG-Ser clade, the CUG-Ser2 clade and the only member of the Alloascoidea clade (*Alloascoidea hylecoeti*) clustering separately from the rest of the clades. This clustering was driven primarily by the codons CUA, CUG, UUG, and UUA (Supplementary Figure 1b), with species in the CUG-Ser, CUG-Ser2 and *A. hylecoeti* being underrepresented in CUA and CUG and overrepresented in UUA and UUG. These four codons are all canonically decoded as leucine, suggesting that the reassignment of the CUG codon in the CUG-Ser1 and CUG-Ser2 clades is largely responsible for the separation of CUG-Ser1 and CUG-Ser2 clades from the rest. This result, however, does not explain the clustering of *A. hylecoeti*, which had the second highest overrepresentation of the UUA codon among the sampled Saccharomycotina, including the CUG-Ser1 and CUG-Ser2 clades. *A. hylecoeti* is the only representative genome of the major clade Alloascoideaceae in the dataset, and its genome contains tRNAs that decode all of the leucine codons, except for CUC. Moreover, there is no evidence of alternative codon usage in this species (Muhlhausen et al. 2018). Additional species in this major clade will need to be sequenced to further understand why *A. hylecoeti* is an outlier in the relative usage of the UUA codon.

**Figure 2.**
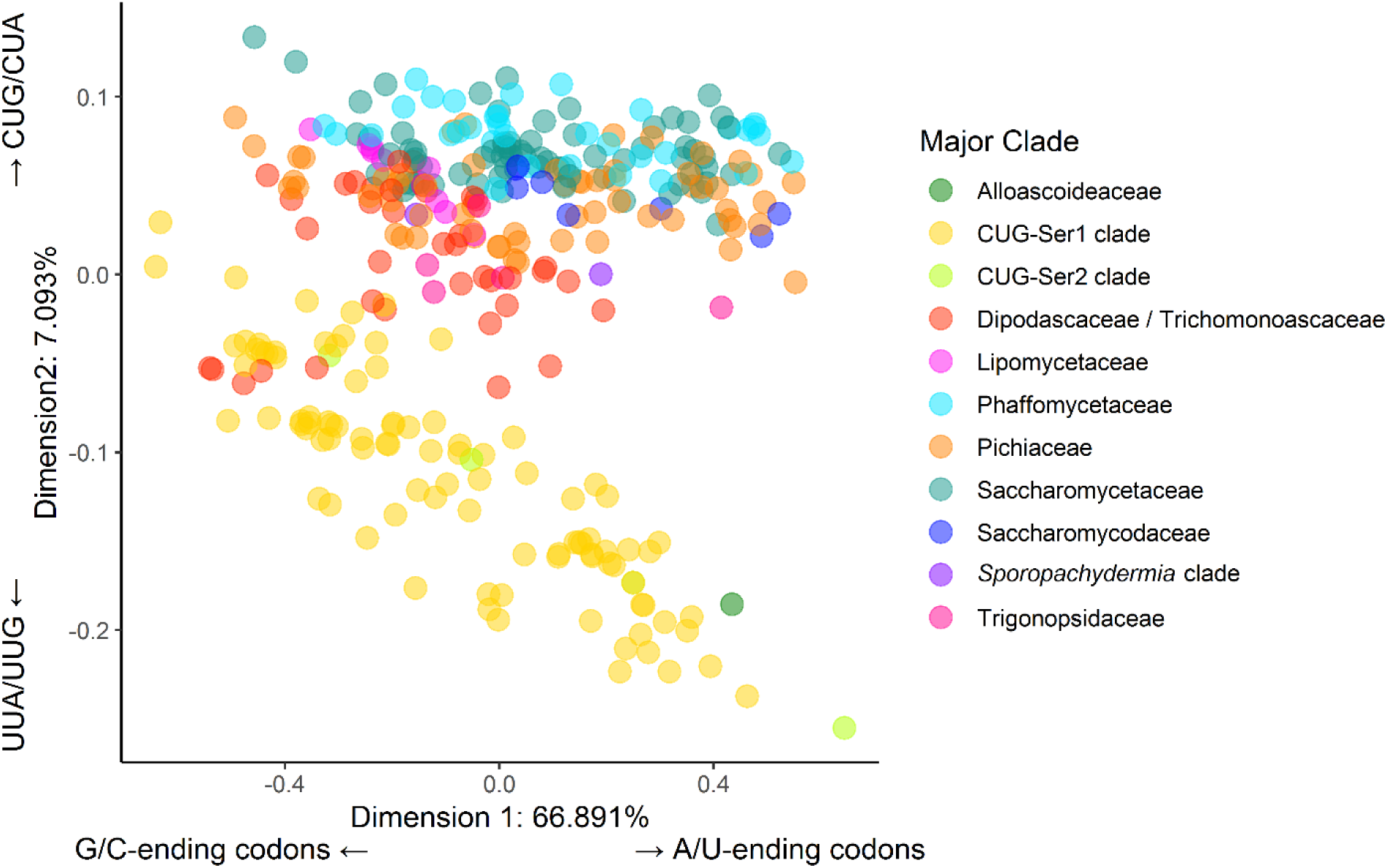
Differences in relative synonymous codon usage values between species are largely driven by variation in the usage of G/C- and A/U-ending codons. The plot shows each of the 327 budding yeast species examined in this study along the first two dimensions (the X and Y axes) of a correspondence analysis. Each axis is labeled with the percent variance explained by the corresponding dimension and the codons that are the major drivers of the observed variance. The first dimension, which explains nearly 67% of the variation between species, is driven by the differential usage of G/C- versus A/U-ending codons. The second dimension, which differentiates the CUG-Ser1 clade, the CUG-Ser2 clade, and one Alloascoideaceae species from the rest of the species in the subphylum, explains a much smaller fraction of the observed variation (about 7%) and is primarily driven by differential usage of the CUA, CUG, UUG, and UUA codons in the two groups.

We next tested whether values of the RSCU metric across species had phylogenetic signal by measuring Pagel’s λ (Pagel 1999) and Blomberg’s K (Blomberg et al. 2003; Ives et al. 2007; Revell 2012) (Supplementary Table 5). Pagel’s λ tests for the presence of phylogenetic signal in a given trait using tree transformation—making the tree more or less star-like. Values for Pagel’s λ vary from 0, which denotes that the trait absence of any phylogenetic signal, to 1, which denotes that the trait varies according to a Brownian model of random genetic drift. Codons’ values for Pagel’s λ ranged from 0.953 (for CUU) to 1 (for multiple codons) with p-values of ≪0.001. These data suggest that codon usage between closely related species is more similar than expected under a Brownian motion model. Blomberg’s K measures the ratio of trait variation among species to the contrasts variance. If the trait varies according to a Brownian model of random genetic drift Blomberg’s K will equal 1. Blomberg’s K however can be greater than 1 which indicates that variance in the trait occurs between clades (versus within.) Interestingly, examination of Blomberg’s K identified between-clade variance (K>1) for only the codons CGA, CCA, UUG, and CUA, with the majority of the variance of the remaining codons present within major clades (K<1). Taken together, Pagel’s λ and Blomberg’s K suggest that the phylogenetic signal for most codons resides towards the tips of the phylogeny and explains variation in RSCU between closely related species. Two of the four codons that have phylogenetic signal deeper in the phylogeny (UUG and CUA) canonically encode leucine and were identified as drivers of the second explanatory axis in the correspondence analysis. This result suggests that the phylogenetic correlation between CGA, CCA, UUG and CUA is not restricted to closely related species and represents phylogenetically-driven differences between major clades, whereas the phylogenetic correlation of most other codons is only between closely related species and not between major clades.

### Individual codon usage is driven by neutral and non-neutral forces

The correspondence analysis of RSCU revealed that major differences in codon usage are largely explained by differences in the usage of G/C- and A/U-ending codons (Fig. 2). To determine the influence of neutral mutational bias on the usage of individual codons, we used Pearson’s correlation and phylogenetic generalized least squares (PGLS) to examine the relationship between codon usage and mutational bias. Across all species, the Pearson’s correlation of GC3 and RSCU revealed that all G/C-ending codons and two A/U-ending codons were positively correlated with GC3 (p-value < 0.001 in all cases) (Supplementary Table 6). The two A/U-ending codons that were positively correlated with GC composition bias were CUU and CGA. Interestingly, CGA was one of the codons identified by Blomberg’s K as being phylogenetically differentiated between clades. It is, therefore, not surprising that CGA and CUU are negatively correlated with GC3 in the phylogenetically corrected PGLS analysis (Fig. 3, Supplementary Table 7). In the PGLS analysis all A/U-ending codons are negatively correlated with GC3 and all G/C-ending codons are positively correlated with GC3. These results reveal that there is a strong correlation between mutational bias and codon usage at the genome level.

**Figure 3.**
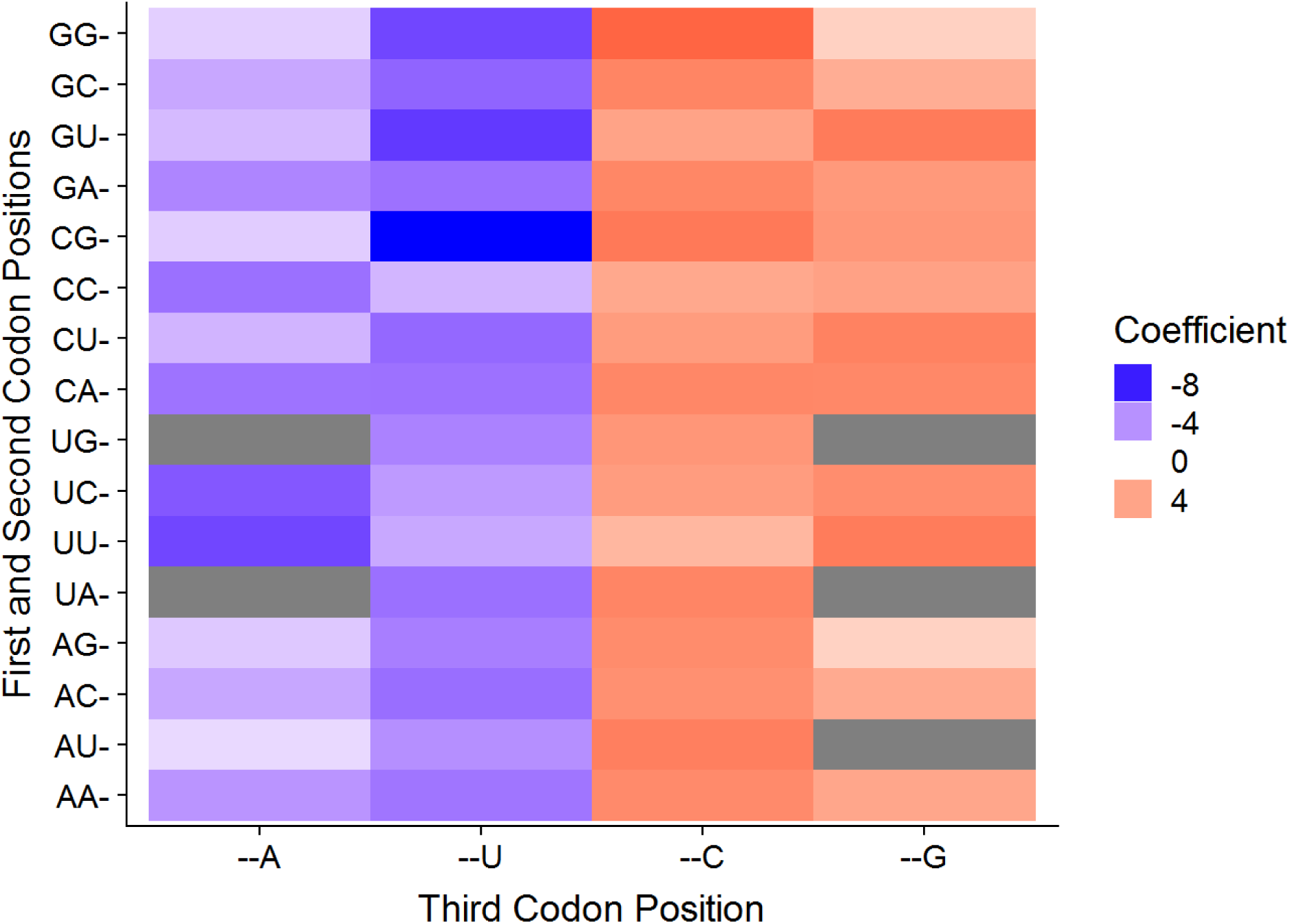
The high correlation between codon usage and GC composition of the third codon position suggests that codon usage bias at the level of individual codons is likely driven by genetic drift. The graph illustrates a phylogenetic generalized least squares comparison between relative synonymous codon usage values and third codon position GC composition (GC3) for each codon across the 327 budding yeast species. Colors toward the red spectrum indicate a positive correlation between CG-ending codons and increasing GC3. Blue colors indicate a negative correlation between A/U-ending codons and increasing GC3. Grey cells denote non-degenerate codons encoding methionine or tryptophan or stop codons.

While the Pearson’s correlation and PGLS analyses suggest that codon bias and GC composition due to mutational bias are correlated, these metrics do not account for the non-linear relationship between GC composition and codon usage. Therefore, we compared observed relative codon frequencies with equilibrium solutions generated by Palidwor et al. (2010). We compared the observed relative codon frequencies for every codon with the equilibrium solutions and measured fit using R^2^ (Fig. 4; Supplementary Table 8). All but one of the 2-fold degenerate codons had an R^2^ value > 0.5 when compared to the neutral expectation (Fig. 4C). For example, the codon GCC fit the neutral expectation very well (R^2^ = 0.671; Fig 4a). The only 2-fold degenerate amino acid encoded by a codon that had an R^2^ < 0.5 was phenylalanine (R^2^ = 0.236). For the 3-fold and 4-fold degenerate codons, the R^2^ values for the individual codons varied but, as previously noted (Palidwor et al. 2010), the summed predictions for G/C-ending codons and A/T-ending codons better fit the neutral expectation (Fig. 4C: second column). The exceptions to this were proline, arginine, and glycine, which showed deviations from the neutral expectation even with the summed statistics (Fig. 4B). To ensure that phylogenetic signal was not driving the deviations from the neutral expectation, we assessed Blomberg’s K of the individual species’ residuals used to compute the R^2^ value. A total of 7 codons had Blomberg’s K variances over 1 (Fig. 4C: Supplementary Table 8), suggesting that deviations from the neutral expectation were driven by differences between major clades. Even after accounting for phylogenetic signal and the improved fit of the summed predictions, codons for proline, glycine, and arginine still showed deviations from the neutral expectation, suggesting that their usages are at least partially driven by selection.

**Figure 4.**
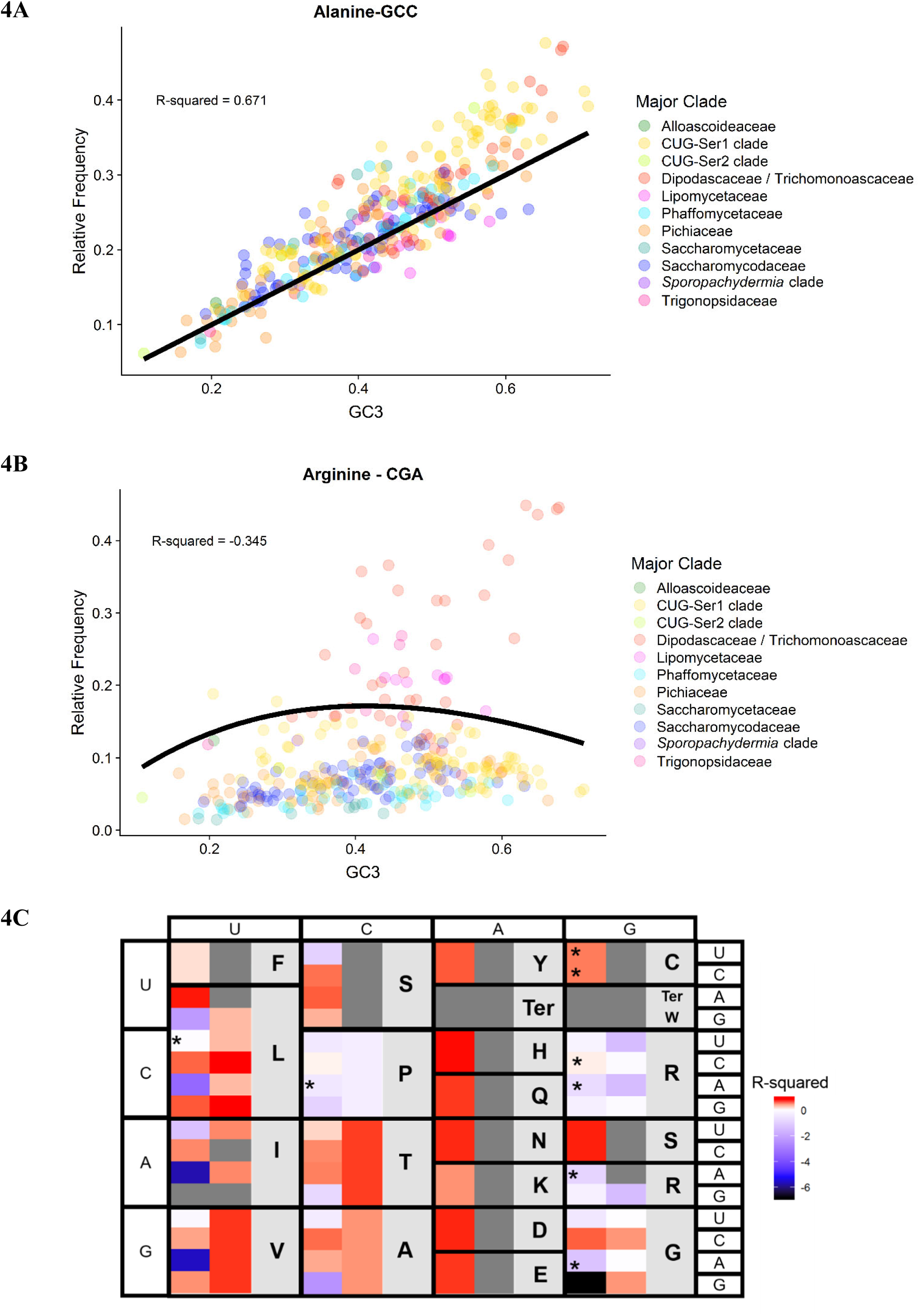
The complex relationship between relative frequency and genome-wide average base composition of the third codon position (GC3) suggests that individual codons vary in their fit to the neutral expectation (i.e., that codon usage is solely driven by GC mutational bias and genetic drift). The neutral expectations for the different codons were obtained from the models developed by Palidwor et al. (2010). A) Observed relative frequency of the alanine codon GCC (shown on the Y axis) plotted against GC3 (shown on the X axis) for each of the 327 budding yeast species analyzed in this study. The codon GCC had a good fit to the neutral expectation (black line, R-squared value = 0.671). B) Observed relative frequency of the arginine codon CGU plotted against GC3 composition for each species. The codon CGU had a poor fit to the neutral expectation (black line, R-squared value = −0.165); the same trend was also observed in the other Group-2 arginine codons (CGA and AGG). C) R-squared values for each of the codons (first column) and the sum of all codons for an amino acid (second column) compared to their neutral expectations. Boxes colored towards the red spectrum indicate a better fit to the neutral model, while boxes colored towards the blue spectrum indicate a poorer fit (i.e., worse than the mean) to the neutral model. Grey-colored boxes in the first column indicate non-degenerate amino acids or stop codons; grey boxes in the second column indicate codons that either have their own models (e.g., ATC) or have values that stem from the same model (e.g., all amino acids encoded by two codons, such as tyrosine (Y), which is encoded by TAT and TAC). Asterisks indicate codons with a Blomberg’s K variance over 1 when comparing GC3 and relative frequency, suggesting that the GC3 and relative frequency values for these codons are correlated due to phylogeny (i.e., closely related species tend to have more similar GC3 and relative frequency values due to shared ancestry).

### Gene-level codon usage does not fit the neutral expectation

To assess the role of mutational bias across all genes within each genome, we next examined the relationship between the ENC of each gene and its GC3s vis-a-vis the neutral expectation (i.e., the relationship between ENC and GC3s if neutral mutational bias were the only force acting on codon usage). For each genome, we computed the number of genes that fell 10% and 20% of the maximum value outside of the neutral expectation between NC and GC3s (dos Reis et al. 2004). Out of a total of 1,683,203 genes, 381,174 (23%) genes fell outside the 10% threshold and 205,558 (12%) fell outside of the 20% threshold (Fig. 5A; Supplementary Table 9). We also tested each species’ overall fit to the neutral expectation by calculating an R^2^ fit to the neutral expectation (Fig. 5B & 5C). This analysis revealed that 7 genomes had R^2^ values greater than 0.5, suggesting that codon usage in these species can largely be explained by neutral mutational bias. Twelve species had an intermediate R^2^ value between 0.25 and 0.5 (or [0.25 – 0.50]), suggesting that neutral mutational bias is partially responsible for codon usage in most genes in these species. Finally, 72 species had low R^2^ values between 0.00 and 0.25, while the remaining 277 species had values below 0. The species with low and negative R^2^values deviate from the neutral expectation, suggesting that mutational bias is not the sole driving factor of codon bias within these genomes.

**Figure 5.**
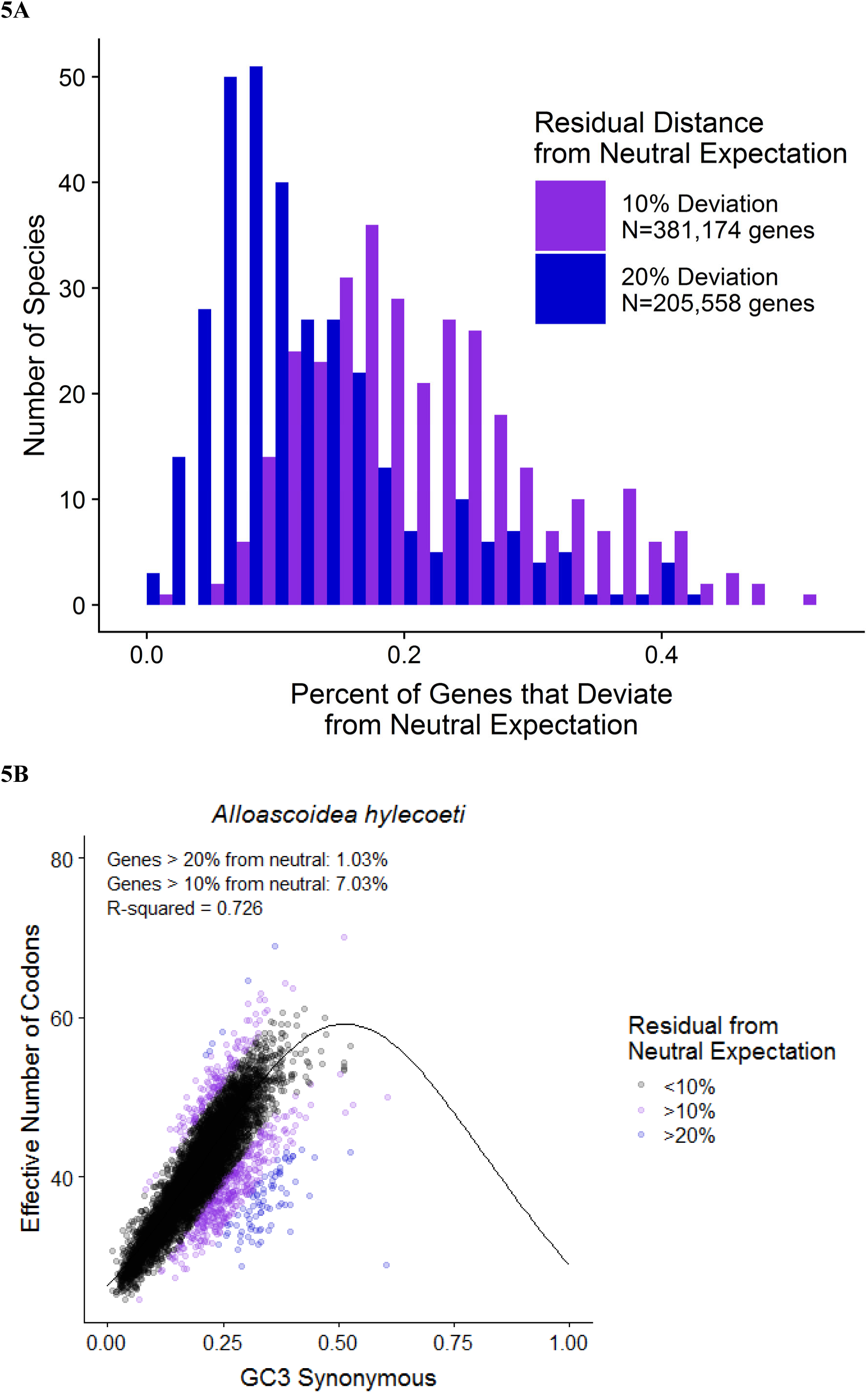
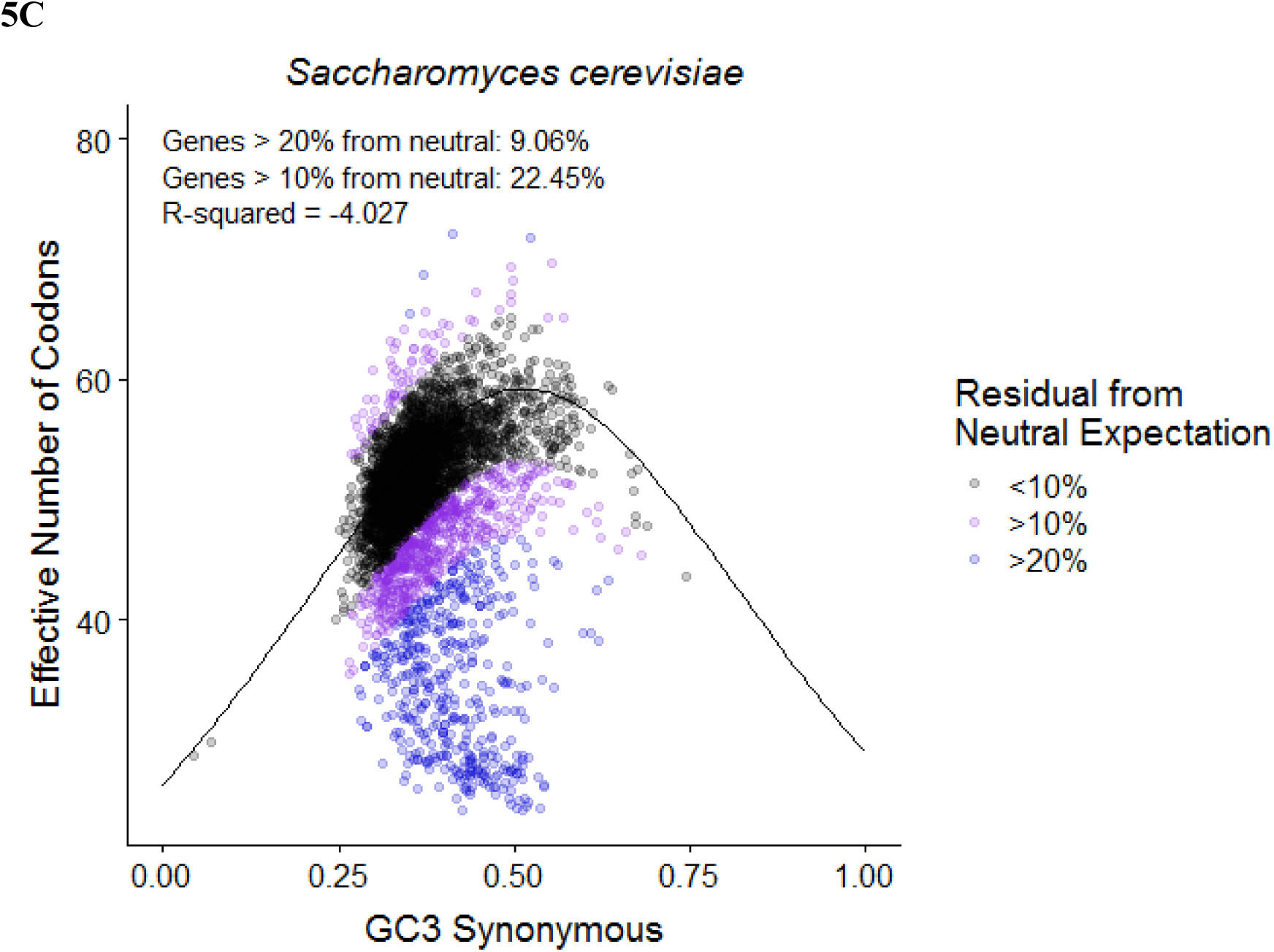
Comparison of the silent third position GC composition (GC3s) to the effective number of codons (Nc) across 327 budding yeast species shows that a significant portion of the genes in many species’ genomes deviate substantially from the neutral expectation. A) Distribution of the percentage of genes that deviate more than 10% (purple bars) or 20% (blue bars) from the neutral expectation. Almost half of the genomes have 10% or more of their genes deviate at the 20% threshold (159 / 327), and almost all of the genomes do so at the 10% threshold (309 / 327). B) The genome of the yeast *Alloascoidea hylecoeti* shows a high correlation between GC3s and Nc (R-squared value = 0.762), in line with neutral expectations. The neutral expectation (i.e., the expectation when the only influence is GC mutational bias and genetic drift) of the effective number of codons for a given GC content of third positions in a genome is indicated by the black line. C) In contrast, the genome of *Saccharomyces cerevisiae* shows a lack of correlation between GC3s and Nc (R-squared value = −4.027) and does not conform with the neutral expectation.

### Codon usage in most budding yeast genomes is under translational selection

The previous analysis suggested that most Saccharomycotina species deviate from the strictly neutral expectation between GC3s and NC within their genomes (Fig. 5). To test whether translational selection influenced codon usage in budding yeast genomes, we calculated the S-value or the amount of selection on codon usage due to tRNA adaptation. To determine the effect of not accounting for CUG codon reassignment in our analysis, we calculated S-values for genomes with CUG and with all CUG codons removed (Supplementary Table 10). The R^2^ value when comparing the S-value for the CUG and CUG-removed datasets was 0.99. This suggests that our results are valid despite not accounting for the codon reassignment. S-values could not be produced for the species *Martiniozyma abiesophila, Nadsonia fulvescens* var. *fulvescens*, and *Botryozyma nematodophila*, because they did not produce viable wi values from stAI-calc due to software issues (Supplementary Table 11). S-values were computed for the remaining 324 species, and significance was assessed using a permutation test (Fig. 6A). Thirty-four species from 6 of the 9 clades did not have S-values that were significant at the 0.05 or 0.95 level in the permutation test (Supplementary Table 10). These non-significant results ranged in S-value between −0.252 and 0.577, with a median value of 0.273. This result suggests that, in these species, gene-level codon usage could not be distinguished from neutral mutational bias; therefore, it is unlikely that translational selection is broadly acting in these species. In contrast, 27 species exhibit moderate S-values between 0.28 and 0.5 (Fig. 6B), on par with levels of translational selection observed in *C. elegans* (S-value of 0.45; dos Reis et al. 2004). A moderately high S-value between 0.5 and 0.75 was observed in 157 species. Finally, a very high S-value above 0.75 was observed for 107 species, including *S. cerevisiae* (Fig. 6C), as previously reported (dos Reis et al. 2004). Overall, 291 / 324 (94%) of genomes examined showed moderate to very high S-values, suggesting that translational selection is widespread across budding yeast genomes.

**Figure 6.**
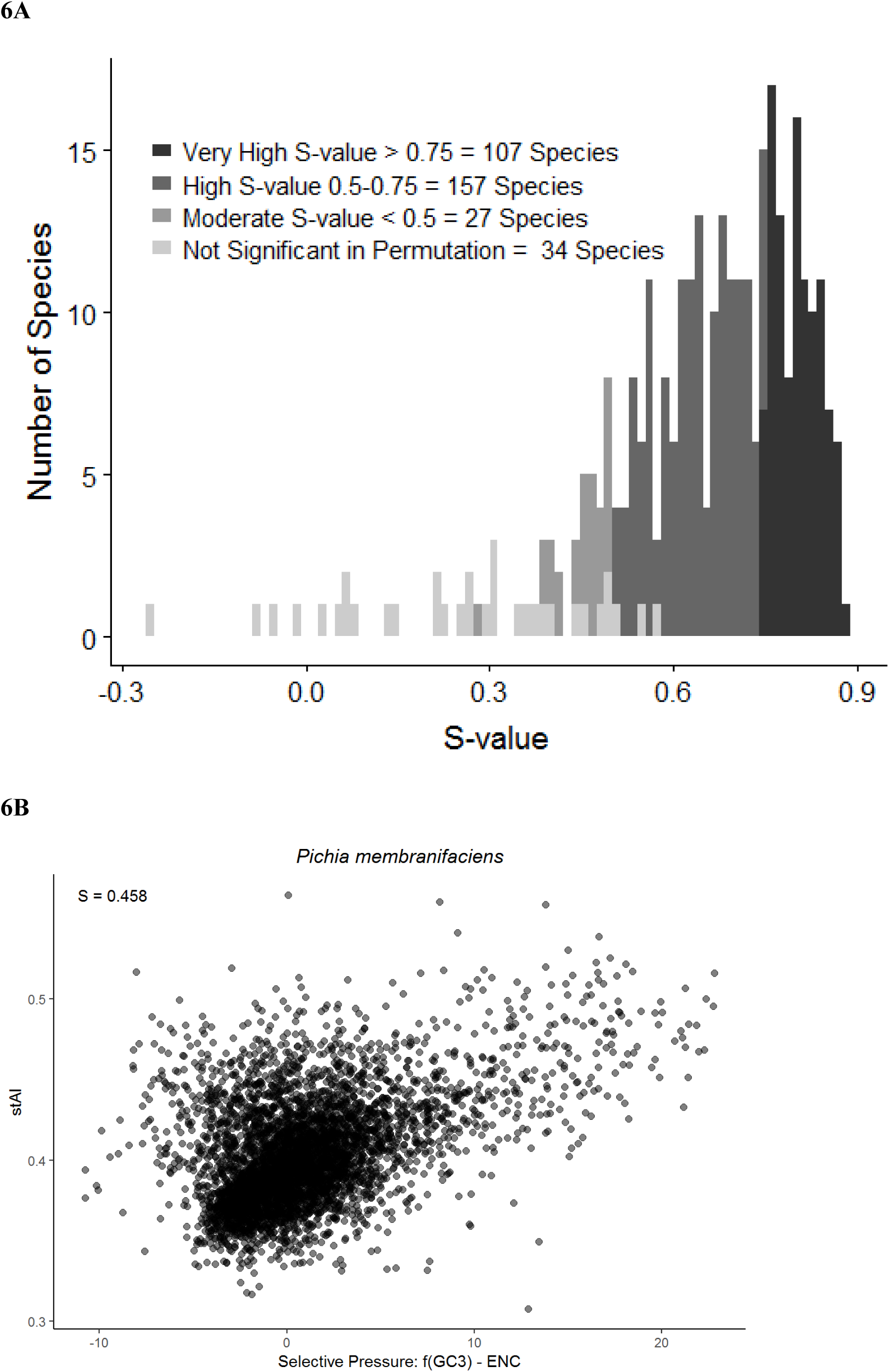
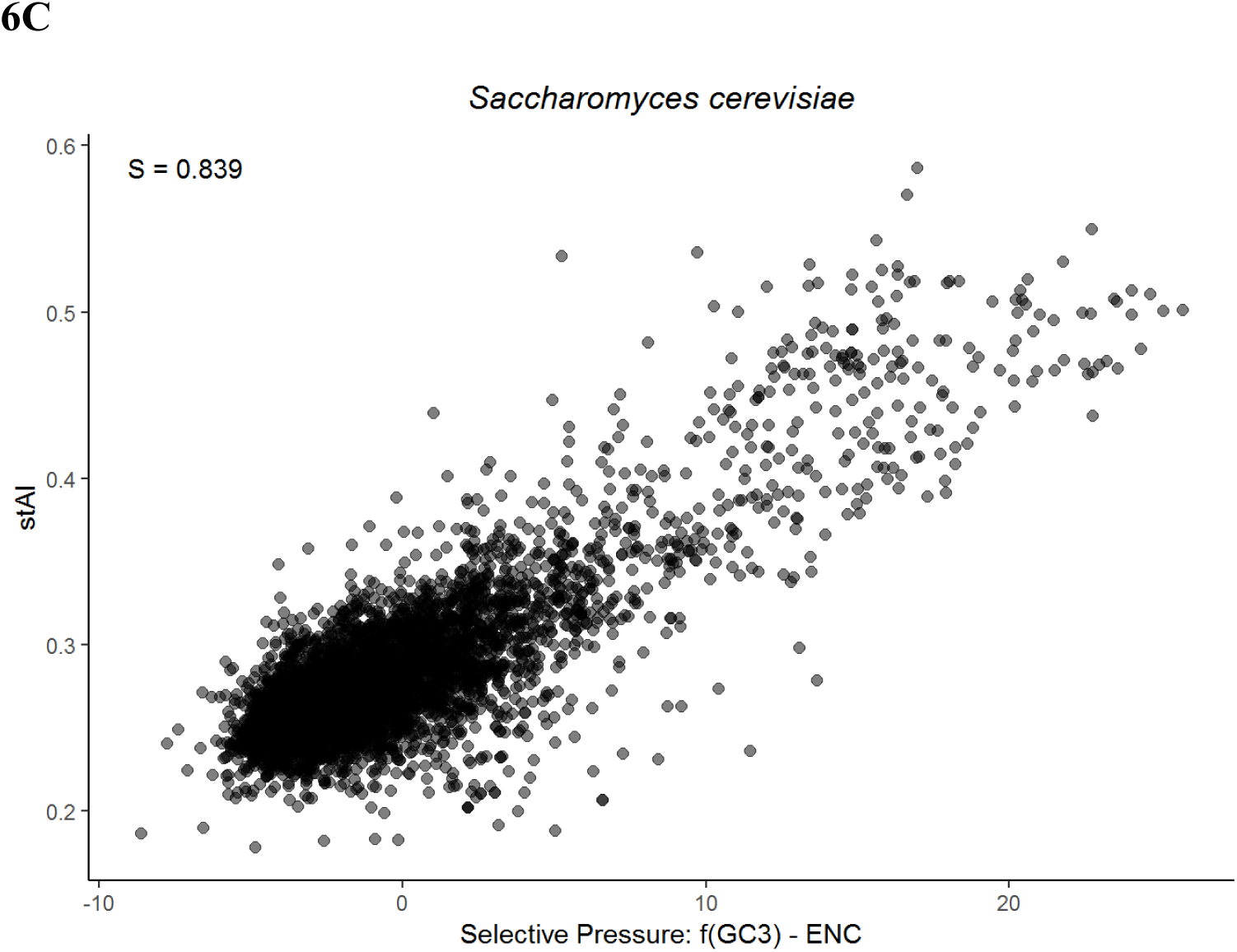
Most genomes in the budding yeast subphylum exhibit moderate to high levels of translational selection on codon bias. Translational selection on codon bias was measured using the S-test, which examines the correlation between the stAI value and the selective pressure (estimated by f(GC3)-ENC where f(GC3) is a modified function of Wright’s neutral relationship between the silent GC content of a gene and the effective number of codons) on all coding sequences in a genome. Each point in the comparison between stAI and selective pressure is a single coding sequence in one genome. Higher S-values indicate higher levels of translational selection on codon bias. A) Distribution of the significant S-values (p<0.05 in permutation test; 293 species out of 327) and non-significant S-values (p>0.05 in permutation test; 34 / 327 species). B) *Pichia membranifaciens*, an example of a species that exhibits low translational selection on codon bias (p<0.05 in permutation test; n=10,000). C) *Saccharomyces cerevisiae*, an example of a species that exhibits high translational selection on codon bias (p < 0.01 in permutation test; n=10,000).

### Translational selection is weakly associated with tRNAome size

To determine which features are associated with S-values, we examined the relationship between S-values with the combinations of two or more of the following features: genome size, tRNAome size, gene number, number of metabolic traits, and number of isolation environments (Supplementary Table 12). The linear model with the highest explanatory power, which accounted for 17.47% of the variation in S-value, includes genome size, tRNAome size, gene number, and total metabolic traits (Supplementary Table 13). Among the four features in the model, tRNAome size had the biggest contribution, followed by genome size, gene number, and reported metabolic traits (0.612 versus 0.229, 0.119, and 0.039, respectively.) To gain further insight into the contribution of the tRNAome size, we tested a Gaussian model (Fig. 7) based on previously reported analyses (dos Reis et al. 2004). The R^2^ value of the Gaussian model was higher than that of the linear model (0.11 vs 0.04), although neither model had a very good fit. The Gaussian model suggests that the maximum selection occurs at an intermediate tRNAome size. Interestingly, the estimated maximum for S-value occurs at a tRNAome size of 336 tRNA genes, a value similar to the tRNAome size that corresponds with the maximum modeled S-value from previous models (tRNAome of about 300) (dos Reis et al. 2004). The phylogenetically corrected PGLS analysis revealed no correlation between S-value and either genome size or tRNAome (Supplementary Fig. 2). Overall, none of the features we tested had strong associations, individually or additively, with S-value, even when phylogenetically corrected.

**Figure 7.**
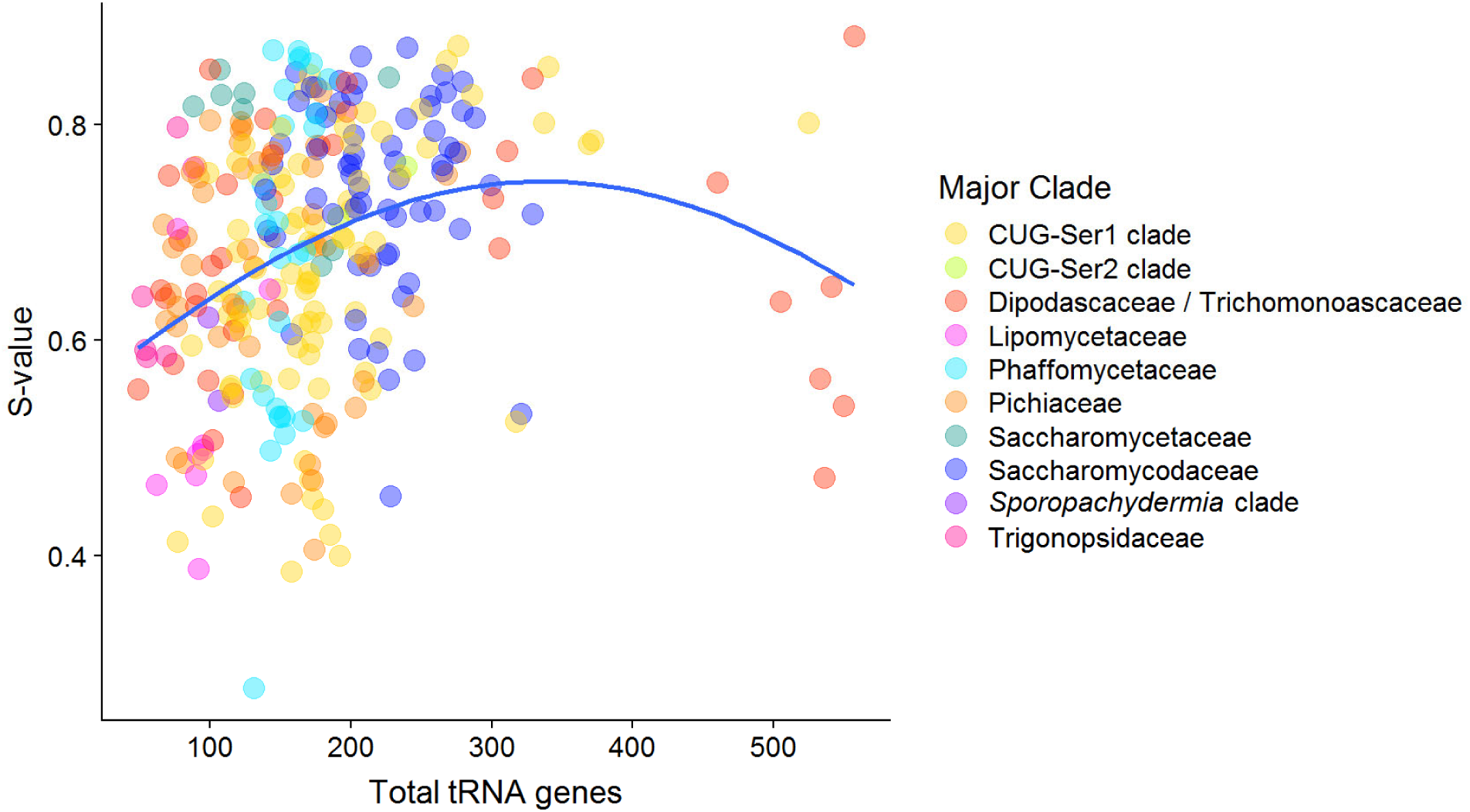
Maximum translational selection occurs at an intermediate number of total tRNA genes in the genome. This plot shows the relationship between the total number of tRNA genes in a genome (tRNAome size) and S-value for each the 327 budding yeast species analyzed in this study. The best fitting model (blue) was a Gaussian distribution with a maximum S-value at 336 tRNA genes. This suggests that species with either low or high numbers of total tRNA genes exhibit lower levels of translational selection.

## Discussion

In this study, we surveyed the patterns and forces underlying codon bias across 327 budding yeasts from the subphylum Saccharomycotina. Cluster, correspondence, and correlation analyses of the relative synonymous codon usage across the subphylum is consistent with mutational bias as a significant driver of codon bias—A/U ending codons are generally overrepresented and G/C ending codons are generally underrepresented. This finding is consistent with the low GC content (average silent GC context of 42%) found across the subphylum. Several previous studies have suggested that genome-wide mutational processes are the primary drivers of genome-wide codon usage (Knight et al. 2001; Chen et al. 2004; Wan et al. 2004), and we clearly observed the influence of these neutral processes at the genome level. Notably, we also found evidence of selection in both specific codons and genes, which we discuss below.

At the level of individual codon usage, two codons in particular—CGA and CUA—had multiple lines of evidence for violating assumptions of neutral GC-mutational bias. For CGA, our results are consistent with previous reports that decoding of the CGA codon in *S. cerevisiae* is inhibitory to translation due to codon-anticodon interactions (Letzring et al. 2010; Letzring et al. 2013). This effect, however, may not be universal across the Saccharomycotina: CGA was underrepresented (RSCU < 1) in 222 species but overrepresented (RSCU > 1) in 105 species. RSCU of CGA also varies between major clades of the Saccharomycotina with the Dipodascaceae/Trichomonascaceae clade having the highest average RSCU (1.47) and the Phaffomycetaceae clade having the lowest average RSCU (0.66). Given that Dipodascaceae/Trichomonascaceae clade is distantly related to Saccharomycetaceae, the major clade that *S. cerevisiae* belongs to, it is likely that the two independent defects in translation that result in the inhibitory nature of CGA in *S. cerevisiae* (Letzring et al. 2013) evolved within Saccharomycetaceae, after the divergence of the two clades. The codon CGA is not the only arginine encoding codon to violate the neutral assumptions (Fig. 4C). Deviations in the remaining arginine codons may be a result of strong directional selection due to the large number of degenerate codons encoding arginine, which may result in more opportunities for poor codon-tRNA pairing (Duret and Mouchiroud 1999; McVean and Vieira 2001).

For CUA, departure from assumptions of neutral GC-mutational bias are likely driven by the reassignment of CUG in the CUG-Ser1 and CUG-Ser2 clades, which had profound effects on the remaining leucine codons since the majority of CUG codons that remained leucine were reassigned to UUG or UUA (Massey et al. 2003; Miranda et al. 2006). This conclusion is supported by the observation that the CUA codon is underrepresented in the CUG-Ser1 and CUG-Ser2 clades (Fig. 1; Supplementary Table 14) compared to other major clades in the subphylum (Fig. 1: Supplementary Table 14). Underrepresentation of CUA is not exclusive to the CUG-Ser2 and CUG-Ser1 clades—the Dipodascaceae/Trichomonascaceae major clade had an average RSCU of 0.60 and includes 12 species (of 37) with a very low RSCU less than 0.5. This may suggest that the Dipodascaceae/Trichomonascaceae major clade experienced similar evolutionary pressures to those that may have contributed to codon reassignment, such as the hypothesized presence of a Virus-Like Element with killer activity in the CUG-Ser1 and CUG-Ser2 clades (Krassowski et al. 2018). The most studied member of the Dipodascaceae/Trichomonascaceae major clade, *Yarrowia lipolytica*, possesses virus-like particles, but these particles do not appear to be associated with a killer phenotype (Tréton et al. 1985; el-Sherbeini et al. 1987). This finding highlights the strong impact of codon reassignment on codon usage.

We also observed deviations from the neutral expectation in all codons that encode proline. Biases in proline codon usage may be related to proline-induced stalling in translation (Artieri and Fraser 2014). This stalling was observed in *S. cerevisiae* riboprofiling data (Artieri and Fraser 2014) and may be related to the slow incorporation of proline into the growing amino acid chain due to its imino side-chain (Pavlov et al. 2009; Doerfel et al. 2013). Additionally, in *S. cerevisiae*, codons for proline and glycine (which also deviate from the neutral expectation) are involved in frameshift suppression via suppressor tRNAs that contain four-base anticodon sequences that allow for frameshift read-through (Donahue et al. 1981; Gaber and Culbertson 1982). As a whole, the results of the codon-specific analysis suggest that while many codons are highly correlated with mutational bias, specific codons may be under a variety of selective forces—especially translational selection—that alter codon usage.

Almost a quarter of the 1,683,203 genes found in the 327 budding yeast genomes deviate from the neutral expectation by at least 10%. These results are consistent with the observation that codon bias varies between transcripts within a species (Sharp et al. 1988; Chen et al. 2004) and is associated with increased expression. In fact, for the species *Saccharomyces mikatae*, the degree to which a transcript differs from the neutral expectation (greater residual) is moderately associated with greater expression at steady state (R2 of 0.414; Supplementary Figure 3; Tsankov et al. 2010). For the majority of the species examined (320), mutational bias is not the only force influencing codon bias among transcripts.

Assessing how translational selection may influence codon usage bias within species, we found that the majority of species exhibited moderate or high contribution of selection to the variation in codon bias (Fig. 6A). Previous work suggested a model in which the highest amount of selection on synonymous codon usage occurs at intermediate genome size. At the lower end of genome size, low selection is hypothesized to be due to the correlation between small genomes and small tRNAomes with low tRNA gene redundancy. In turn, low tRNA gene redundancy restricts the ability of selection to act on codon bias (Kanaya et al. 1999; dos Reis et al. 2004). At the larger end of genome size, low selection is hypothesized to be due to drift in species with small effective population sizes: this drift would increase the genome size and decrease the ability of selection to shape codon usage (Bulmer 1991). Within Saccharomycotina, the role of tRNAome size is consistent with these predictions, except for genome size. This exception is likely due to a low correlation between genome size and tRNAome size in this group. While tRNAome size and genome size are positively correlated when analyzed using a phylogenetically independent contrast (PIC) (Felsenstein 1985), this correlation is not very strong (adjusted R^2^ of 0.1629.)

In summary, we find that the balance between neutral and selective forces on codon usage varies between genomes, between codons, and between genes within a genome. Some Saccharomycotina species exhibit nearly neutral codon usage in line with those observed in humans or bacteria, such as *Helicobacter pylori*, while other budding yeast species show extremely high adaptation to the tRNA pool through translational selection (dos Reis et al. 2004). This range in the magnitude of forces acting on codon usage in the Saccharomycotina and the low explanatory power of the factors examined suggest that it is difficult to predict *a priori* selection on codon bias based on lineage, cellularity, genome size, tRNAome, or GC composition.

There is moderate to strong evidence for translational selection in most budding yeast genomes examined. This trend may be due to the rapid growth that characterizes most budding yeasts: growth efficiency has been linked to translational selection in codon usage (Andersson and Kurland 1991; Kurland 1991). One interesting implication of this abundance of translational selection is that codon optimization may be a useful proxy for highly expressed genes. It has long been known that ribosomal genes are among both the most highly expressed and highly codon usage-optimized genes across species (Shields et al. 1988; Sharp et al. 1995), leading to their use as the basis for the codon adaptation index (Sharp and Li 1987; Nakamura and Tabata 1997). In our dataset, there are 11,047 genes (average of 35 per species) that are as highly or more highly optimized than the ribosomal genes, suggesting there is a wealth of information about which genes may be highly expressed or differentially highly expressed across this lineage.

## Acknowledgements

We thank the members of the Rokas and Hittinger labs, in particular Xing-Xing Shen, for their feedback and discussions on this project. We would also like to thank the other members of the Y1000+ project (http://www.y1000plus.org/) including, Jacek Kominek and Xiaofan Zhou, for their feedback. We would also like to thank Renana Sabi, Renana Volvovitch Daniel and Tamir Tuller, the creators of stAIcalc, for their assistance in troubleshooting the codon adaptation analysis.

